# The Influences of Stomatal Size and Density on Rice Drought, Salinity and VPD Resilience

**DOI:** 10.1101/2022.05.13.491841

**Authors:** Robert S. Caine, Emily L. Harrison, Jennifer M. Sloan, Paulina M. Flis, Sina Fischer, Nguyen Trong Phuoc, Nguyen Thi Lang, Julie E. Gray, Holly L. Croft

## Abstract

- A warming climate coupled with reductions in water availability and rising salinity are increasingly affecting rice yields (*Oryza sativa* L.). Elevated temperatures are causing vapour pressure deficit (VPD) rises, leading to stomata closure, further reducing plant productivity and cooling. It is unclear which conformation of stomatal size (SS) and stomatal density (SD) will best suit these future environmental extremes.
- To understand the influence of stomatal characteristics on rice abiotic stress tolerance, we screened the stomatal characteristics of 72 traditionally-bred varieties. We found significant variation in SS, SD and maximal stomatal conductance (*g_smax_*) but did not identify any varieties with SD and *g_smax_* as low as the genetically manipulated stomatal development mutant *OsEPF1oe*.
- Traditionally-bred varieties with high SD and small SS (resulting in high *g_smax_*) typically had lower biomasses, and these plants were more resilient to drought than low SD and large SS plants, which were physically larger. None of the varieties tested were as resilient to drought or salinity as low SD *OsEPF1oe* mutants. High SD and small SS rice displayed faster stomatal closure during rising VPD, but photosynthesis and plant cooling were reduced.
- Compromises will be required when choosing rice SS and SD to tackle multiple future environmental stresses.

## Introduction

Developing high yielding rice varieties that use less water which can withstand multiple abiotic stresses will be critical for maintaining future global food security (Singh *et al*., 2021). Currently, rice is the most consumed human food crop, providing more than 20% of the total calories consumed worldwide (Fukagawa & Ziska, 2019). It takes around 2500 litres of water to produce 1kg of grain, and globally, this equates to around 30% of all the world’s fresh water supplies being used for rice production (Bouman, 2009). Half of all cultivation is irrigated, and this boosts crop yields and protects against drought and heat stress, but such practices are highly water-intensive often leading to anaerobic soils and excessive methane production (Singh *et al*., 2021). The majority of other farmed rice is rain fed, with 34% grown in lowland ecosystems, and 9% in upland environments. As the human population continues to grow, and as climate change intensifies, clean water for irrigation will decrease, and this combined with a need to reduce emissions will prompt a move towards more rain fed rice agriculture (Bouman, 2009; Singh *et al*., 2021). Such changes in farming practices are forecast to occur at the same time as increased incidences of severe droughts, higher temperatures and rising soil salinity (Livsey *et al*., 2019; IPCC, In press.). Taken together these factors have the potential to massively impact global rice yields exactly when demand for rice (and clean water) is rapidly growing (Panda *et al*., 2021; Singh *et al*., 2021).

Plant water-use is controlled by stomata – microscopic epidermal pores comprised of a pair of guard cells that regulate CO_2_ uptake for plant photosynthesis (*A*), with the concurrent release of water via alterations to stomatal conductance (*g_sw_*). As well as regulating gaseous exchanges, stomata facilitate water and nutrient uptake from soils and also aid in plant cooling via increases to the transpiration (*E*) stream when temperatures rise. Under drought, salinity or increasing vapour-pressure deficit (VPD), stomatal apertures reduce or close, and this restricts photosynthesis, water loss, nutrient uptake, plant cooling and ultimately growth and seed yield (Merilo *et al*., 2018; Grossiord *et al*., 2020; Ma *et al*., 2020). Rises in VPD occur when the difference between the maximum amount of water the air can hold, and the actual amount of water in the air increases. This is often the case as temperatures rise (Grossiord *et al*., 2020), with forecasts predicting VPD will continue to rise until the end of the century (Yuan *et al*., 2019). Such rises in VPD have thus far offset any CO_2_ fertilisation effects associated with rising CO_2_ concentration.

With prolonged changes in environmental stimuli, many plant species are able to modulate their stomatal development by altering stomatal size (SS) and/or stomatal density (SD) – often in opposite directions (Franks & Beerling, 2009; Franks *et al*., 2012). This developmental adjustment has been widely observed in living plants, herbarium samples and even fossil records, and coincides with CO_2_ fluctuations that have occurred during different geological epochs. Typically, high CO_2_ environments are associated with an increased SS and reduced SD, and low CO_2_ concentrations are associated with the opposite conformation of SS and SD (Franks & Beerling, 2009). Such developmental adjustments have been suggested to alter *g_sw_* range by permitting a higher calculated anatomical maximum stomatal conductance (*g_smax_*); with plants that have a high SD and small SS being able to potentially achieve high operating *g_sw_* and *g_smax_* levels (Franks & Beerling, 2009; Bertolino *et al*., 2019). Alterations to SS and SD have also been suggested to adjust the speed at which stomata respond to environmental cues; with evidence suggesting that plants with smaller SS and higher SD are more rapidly able to respond to environmental conditions, and this can boost water-use efficiency (McAusland *et al*., 2016; Bertolino *et al*., 2019). For fluctuating light responses, this means faster responsivity of *g_sw_* to changes in *A*, which reduces unnecessary water loss (Lawson & Vialet-Chabrand, 2019). Examples of negative correlations between SS and speed have been observed within a species (or closely related species) but not necessarily between distantly related species (McAusland *et al*., 2016). However, other studies have shown that stomatal responsiveness is not always related to SS (Eyland *et al*., 2021), and the improved benefits of small SS on optimising *A* may be light dependent (Zhang *et al*., 2019). In monocot grasses such as rice, each stomatal guard cell pair is surrounded by a pair of subsidiary cells, which increase the speed of stomatal opening and closure (Franks & Farquhar, 2007; Raissig *et al*., 2017; Gray *et al*., 2020).

It is possible to improve various abiotic stress responses by genetically manipulating stomatal physiology or stomatal development (Huang *et al*., 2009; Mohammed *et al*., 2019). Previous work manipulating the levels of Epidermal Patterning Factor (EPF) signalling peptides in several species, has shown that SD reductions of up to approximately 50% can lead to significantly lower *g_sw_* without significantly impacting *A* (Hepworth *et al*., 2015; Caine *et al*., 2019; Dunn *et al*., 2019; Mohammed *et al*., 2019). These larger reductions in *g_sw_* compared to *A* led to improved intrinsic water-use efficiency (iWUE) without negatively impacting seed yield. In fact, moderate reductions in SD, improved rice yields following drought imposition during the flowering stage (Caine *et al*., 2019). Surprisingly, EPF-driven reductions in SD had inconsistent effects on SS between different rice varieties despite sampling at the same development stage (Caine *et al*., 2019; Mohammed *et al*., 2019). Overexpression of the *OsEPF1* gene in transgenic IR-64 plants previously displayed smaller stomata with lower SD (Caine *et al*., 2019), whereas *OsEPF1oe* Nipponbare plants had larger stomata (Mohammed *et al*., 2019).

It is now clear that reductions in SD can result in reduced crop water loss and increased drought tolerance (Caine *et al*., 2019; Mohammed *et al*., 2019). In this study, we investigate whether selecting for specific SD and SS traits could mitigate against not only drought, but also additional climate change associated abiotic stresses including rising salinity and VPD. Specifically, we ask: 1) Is it possible to identify the combination of SS and SD found in IR-64 *OsEPF1oe* (*OsEPF1oe*) in other traditionally-bred high yielding rice varieties? 2) Do *OsEPF1oe* or other plants with fewer (or smaller) stomata perform better under drought or saline stress conditions? And 3) does SD and/or SS affect stomatal responses to high temperature and increased VPD.

## Material and Methods

### Plant Materials

A collection of 72 rice varieties previously assayed for salinity tolerance were kindly provided by Jose De Vega, Earlham Institute (Table. S1). Two independently transformed lines of IR-64 variety overexpressing *OsEPF1* have been previously described (Caine *et al*., 2019).

### Plant growth conditions

Rice seeds placed in 15-20 ml RO water were sealed in Petri dishes with micropore tape (3M, Saint Paul, Minnesota) and germinated under: 12 h 26°C: 12 h 24°C light:dark cycle at photosynthetically active radiation (PAR) 200 μmol m^-2^ s^-1^. Seedlings were sown onto a previously described soil mix (Caine *et al*., 2019), in 0.8 L pots (IPP, Bytom, Poland). Pots were prepared by first half filling with soil mix, then RO water was mixed through, and then a second equal application of soil was added, and further RO water mixed through to saturate the soil. When drained the soil level was c. 1.5 cm from the pot apex. For preparation of salt-treated pots in Fig. 4, a 20 mM solution of NaCl rather than RO water was applied to saturate the soil during mixing. Seedlings were grown in Conviron growth cabinets (Controlled Environments Ltd, Winnipeg, MB, Canada) to 12 h 30°C: 12 h 24°C light: dark cycle, RH 60% with CO_2_ concentration between 450-480 ppm. For plants in Fig. 4 and Fig. S1, PAR was set at canopy level to 1000 μmol m^-2^ s^-1^. For all other experiments, PAR was 1500 μmol m^-2^ s’^1^. For salt-treated plants in Fig. 5, all samples were initially transplanted into fresh-water-mixed soil, with 50 mM NaCl first applied 8 days after transferring (16 DPG). A constant supply of RO water or salt water was available during experiments.

**Fig. 1.**
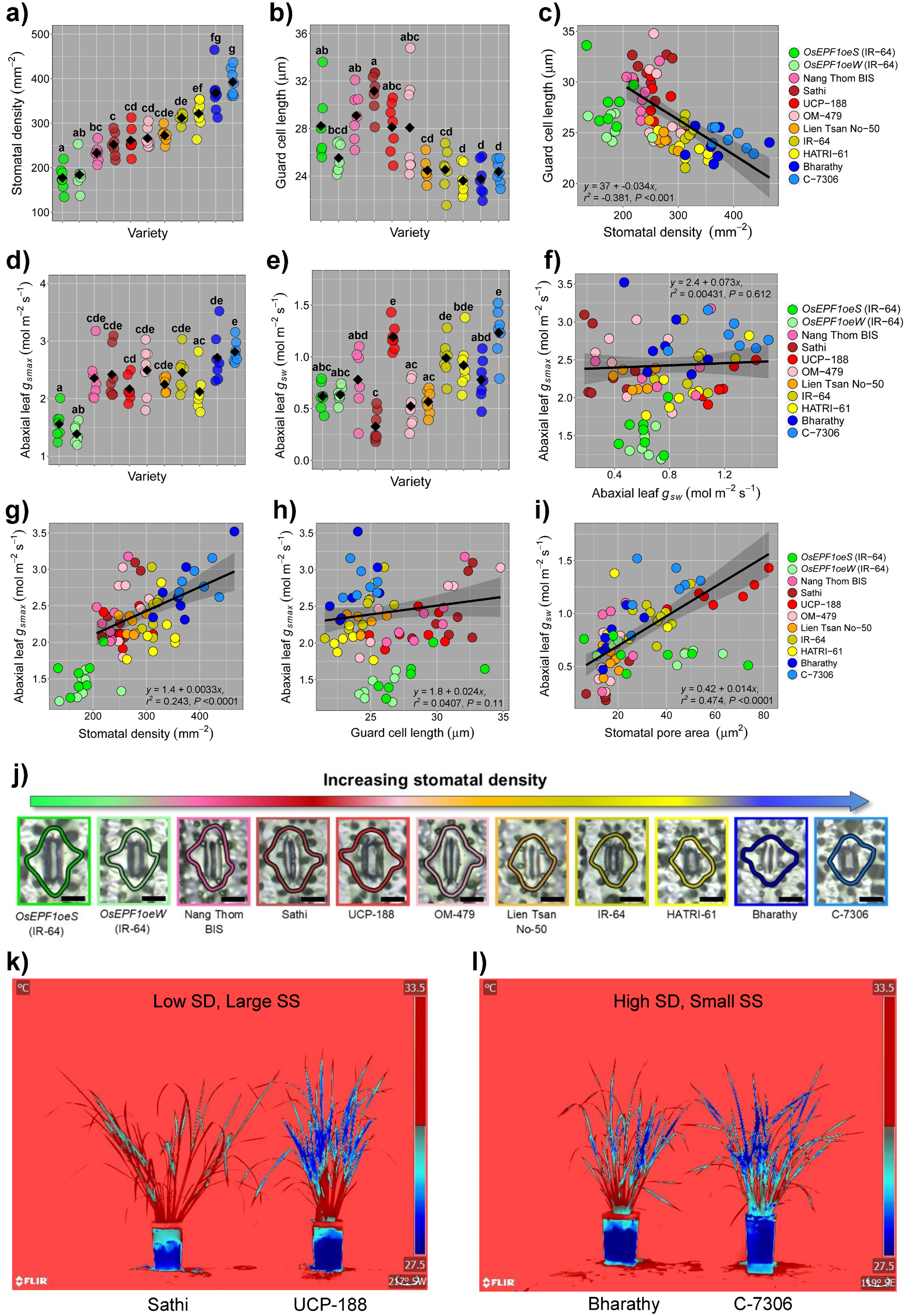
Stomatal size, density and pore aperture contributions to rice gaseous exchange on plants grown in high light conditions (1500 PAR). (a) Abaxial stomatal density (SD) plotted low to high of nine selected rice varieties and two transgenic *OsEPF1oe* plants. (b) Corresponding stomatal size (SS) (guard cell length) measurements of plants in (a). (c) Regression analysis highlighting relationship between SS and SD. (d) Calculated maximal stomatal conductance (*g_smax_*), and (e) corresponding operating stomatal conductance (*g_sw_*) of abaxial leaf surface. (f-i) Regression analysis of (f) abaxial *g_smax_* and operating *g_sw_*, (g) abaxial *g_smax_* and SD, (h) abaxial *g_smax_* and SS and (i) abaxial *g_sw_* and stomatal pore area. (j) Examples of stomatal openness (bars, 10μm). (k and l) Thermal images illustrating temperature differences in plant temperature driven by changes in pore apertures between (k) plants with low SD and large SS and (l) high SD and small SS. Different letters on graphs indicate a significant difference between the means (One-way ANOVA, Tukey HSD test, *P* < 0.05). Black diamonds represent means. Regression analysis and trend lines are based on linear models. *OsEPF1oe* plants are excluded from regression analyses. (a-i) *n* = 7 plants.

### Drought and salinity experiments

For droughted plants (n = 10 or 11), water was withheld for 5 days from 30-35 days post germination (DPG). New leaves were classified as visible new growth emerging from sheaves of pre-existing tillers at 42 DPG. For salinity experiments, salt water was applied when required, and trays changed weekly for 20 mM NaCl experiments (Fig. 4) (n = 6), or two-weekly for the 50 mM NaCl experiment (n = 7 to 9, Fig.5). For both experiments, salt water or fresh water (for controls), was applied from above every time salt trays were changed. Leaf blade and tiller base (2 cm from soil) Φ PSII measurements were collected using a FluorPen FP 110 (PSI, Drasov, Czech Republic). Leaf discs were collected from 35 day-old fresh water and salt grown plants and dried in a 40°C oven for 3 days prior to analysis.

### Gas exchange and thermal imaging measurements

Measurements of *g_sw_* shown in Fig. 1 and 2 were collected using a LI-600 porometer (LI-COR, Lincoln, USA) set to a flow rate of 150 μmol s^-1^ (*n* = 7). Thermal images in Fig.1 were captured using a FLIR T650sc (Wilsonville, USA). Steady state and dynamic gas exchange experiments were performed using LI-6800 Portable Photosynthesis Systems (LI-COR, Lincoln, USA) and attached MultiphaseFlash Fluorometer (6800-01A). Fully expanded leaves of 19-25 day old plants (leaf 5 or 6) were used to collect steady-state measurements in Fig. 4. Leaf chamber conditions were set to light intensity 2000 μmol m^-2^ s^-1^ PAR, relative humidity (RH) 60%, Tair 30°C, flow 300 μmol s^-1^ and [CO_2_]ref 480 ppm. Over a 5-minute period, 10 readings were taken then averaged (*n* = 6 plants). For VPD gas exchange experiments in Fig. 6, measurements were conducted on fully expanded leaf 5 of 19-23 DPG plants, with the leaf chamber set to 2200 μmol m^-2^ s^-1^ PAR, 55% RH, Tair 32°C, flow 400 μmol s^-1^ and 450 ppm [CO_2_]ref. For fluctuations in RH during this experiment see Fig. S3. Once steady-state was reached, two readings were collected at the end of 6-minute intervals, then the temperature was increased by 2.5°C in 4x 6 minute intervals, with readings taken immediately prior to each subsequent temperature increase. Four further readings were recorded at 42 °C (*n* = 7 or 8 plants). Leaf chamber matching was conducted prior to each reading. For steady-state VPD experiments in Fig. S3, chamber settings were the same as in Fig. 6, except Tair was 39°C and RH maintained at 55% throughout (n = 5 or 6).

### Analysis and quantification of stomatal traits

Epidermal imaging and quantification was conducted on IMAGEJ using nail varnish impressions of dental resin imprints taken from leaf 5 (19-23 DPG), (Fig. 4, 6 and Fig. S1 and 3) or leaf 8 (28-33 DPG), (Fig1). Two 0.44 mm^2^ fields of view (FOV) per replicate were used to calculate SD with 5 stomata per biological replicate used to assess guard cell length. Calculations for *g_smax_* were performed as in Caine *et al*. (2019) based on double end-corrected version of the Franks & Farquhar (2001) equation from Dow *et al*. (2014). Graphs and statistical analysis were produced using R software.

### Inductively Coupled Plasma-Mass Spectrometry

Sample preparation was as described previously (Danku *et al*., 2013). In brief, dried plant material was digested with 1ml concentrated nitric acid (trace metal grade, Fisher Chemicals) spiked with Indium (internal standard) in dry block heaters (SCP Science; QMX Laboratories) at 115°C for 4h. The samples were then diluted to 10ml with Milli-Q Direct water (18.2 MΩcm, Merck Millipore) and analysed using ICP-MS (PerkinElmer NexION 2000) in the collision mode (He). Reference material (pooled samples) was run to correct for variation within ICP-MS analysis run. Calibration standards were prepared from single element standards solutions (Inorganic Ventures, Essex Scientific Laboratory Supplies Ltd, Essex, UK). The final element concentrations were obtained by normalizing concentrations to sample dry weight.

## Results

### Rice stomatal size and density are negatively correlated and contribute to operating and maximum potential stomatal conductance

By overexpressing an Epidermal Patterning Factor, we previously showed that reductions in SD reduces rice water requirements and thus improves iWUE and drought tolerance (Caine *et al*., 2019; Mohammed *et al*., 2019). Here we carried out a screen of 72 traditionally-bred rice varieties alongside two independently transformed *OsEPF1oe* lines to survey and compare stomatal traits between genetically engineered plants and a non-transgenic population (**Fig. S1, Table. S1**). Although SD differed widely across the collection, we were unable identify a non-transgenic variety with a mean SD as low as either of the IR-64 *OsEPF1oe* rice lines. We did, however, observe a negative correlation between SS and SD within the full collection of traditionally-bred varieties (*r^2^* = −0.17, *P* < 0.0001, **Fig. S1**), with lower SD varieties typically having larger SS (with the exception of *OsEPF1oe* lines), and higher SD varieties typically having smaller SS. To study how these differences in stomatal traits might affect abiotic stress responses to increased drought, salinity and VPD, we selected nine varieties spanning the range of SS and SD, and conducted a series of experiments alongside *OsEPF1oe* plants. We began by measuring and comparing stomatal morphology, gas exchange and pore size on the leaves of tillering rice plants (**Fig. 1**).

Assessment of SD on leaf 8 of tillering rice revealed traditionally-bred varieties had mean SD values ranging from 233-393 stomata per mm^-2^ whereas the two independent *OsEPF1oe* lines had c. 180 stomata per mm^-2^ (**Fig. 1a**). As expected, a negative correlation between SS and SD was observed between varieties, with those with higher SD typically having smaller guard cells (*r^2^* = −0.38; *P* < 0.0001, **Fig. 1b** and **c**). *OsEPF1oe* lines, which had the lowest SD, did not follow the same trend line, but surprisingly, by the leaf 8 stage (28-33 DPG), stomata were either equal to (*OsEPF1oeW*) or significantly larger (*OsEPFoe1S*) than IR-64 control plants. In the case of *OsEPF1oeS*, this result was opposite to what we had previously found in IR-64 plants at the leaf 5 stage (Caine *et al*., 2019). Despite having larger than expected stomata, very low SD led to *OsEPF1oe* plants having the lowest *g_smax_* (**Fig. 1d**). Within the nine varieties selected, there were limited differences between calculated *g_smax_*, but notably, the varieties with the two highest SD also had the highest mean *g*smax values.

To investigate if operating *g_sw_* followed a similar trend to *g_smax_*, we measured leaf *g_sw_* using a porometer (**Fig. 1e**). Whilst we found no overall correlation between operating *g_sw_* and calculated *g_smax_* across varieties (**Fig. 1f**), we did detect positive relationships between SD and *g_smax_* (*r^2^* = 0.24; *P* < 0.0001, Fig. 1g) and between SD and operating *g_sw_* (*r^2^* = 0.13; *P* < 0.01, **Fig. S2a**). There was no significant correlation between SS and *g_smax_* (*r^2^* = 0.04; *P* = 0.11, **Fig. 1h**), but we did identify a weak negative relationship between SS and *g_sw_* (*r^2^* = 0.09; *P* < 0.02, **Fig. S2b**). Despite very low calculated *g_smax_*, both *OsEPF1oe* lines were able to maintain a similar operating *g_sw_* comparatively to some of the other selected varieties even though SD was around 30% lower (**Fig. 1a, d, e**), and whilst low SD typically meant low operating *g_sw_*, UCP-188 bucked this trend having the equal highest *g_sw_* (**Fig. 1a, e**). Our results suggested that factors other than SS and SD drove the observed differences in operating *g_sw_*, so we next assessed stomatal pore area. Overall we observed a robust correlation (*r^2^* = 0.47; *P* < 0.0001) between gas exchange and the extent of stomatal opening across the nine selected varieties (**Fig. 1i, j**). Rates of stomatal water loss were further explored by assessing whole-plant surface temperatures as a proxy for evaporative transportation – with lower temperatures indicative of higher water loss. Thermal imaging confirmed firstly that the open-pored UCP-188 variety with large stomata was cooler than the Sathi variety with equivalent SS and SD, and secondly, that the open-pored C-7306 variety, with high SD was cooler than the Bharathy variety which again had similar conformations of SS and SD. These observed similarities between surface temperatures and leaf *g_sw_* values, indicate that in addition to SS and SD, stomatal pore aperture (and perhaps other root and vascular features) can differ substantially between rice varieties and this also has the potential to greatly influence water loss (**Fig. 1i-l**).

To investigate if the differences in stomatal morphology and physiology (**Fig. 1**) were associated with overall plant growth, we next measured vegetative-stage above ground biomass at 35 DPG (**Fig. 2**). Within the nine selected varieties, we found that low SD varieties typically had greater biomass than plants with higher SD (**Fig. 2a**). The exceptions to this were the *OsEPF1oe* transgenic plants. With *OsEPF1oe* transgenic lines excluded from regression analyses, a moderate negative correlation was observed between SD and plant biomass (*r^2^* = −0.24; *P* < 0.001), and a weaker positive relationship was detected between SS and biomass (*r^2^* = 0.16; *P* < 0.01) (**Fig. 2b-c**). We also found that pore area was weakly associated with greater plant biomass (*r^2^* = 0.09; *P* < 0.02) (**Fig. 2d**), but no such relationship was detected for operating *g_sw_* (**Fig. 2e**). Assessment of *g_smax_* values revealed a weak negative correlation with plant biomass (*r^2^* = −0.07; *P* < 0.04) (**Fig. 2f**), with smaller plants typically having marginally higher *g_smax_* values, although this was not the case for *OsEPF1oe* plants.

**Fig. 2.**
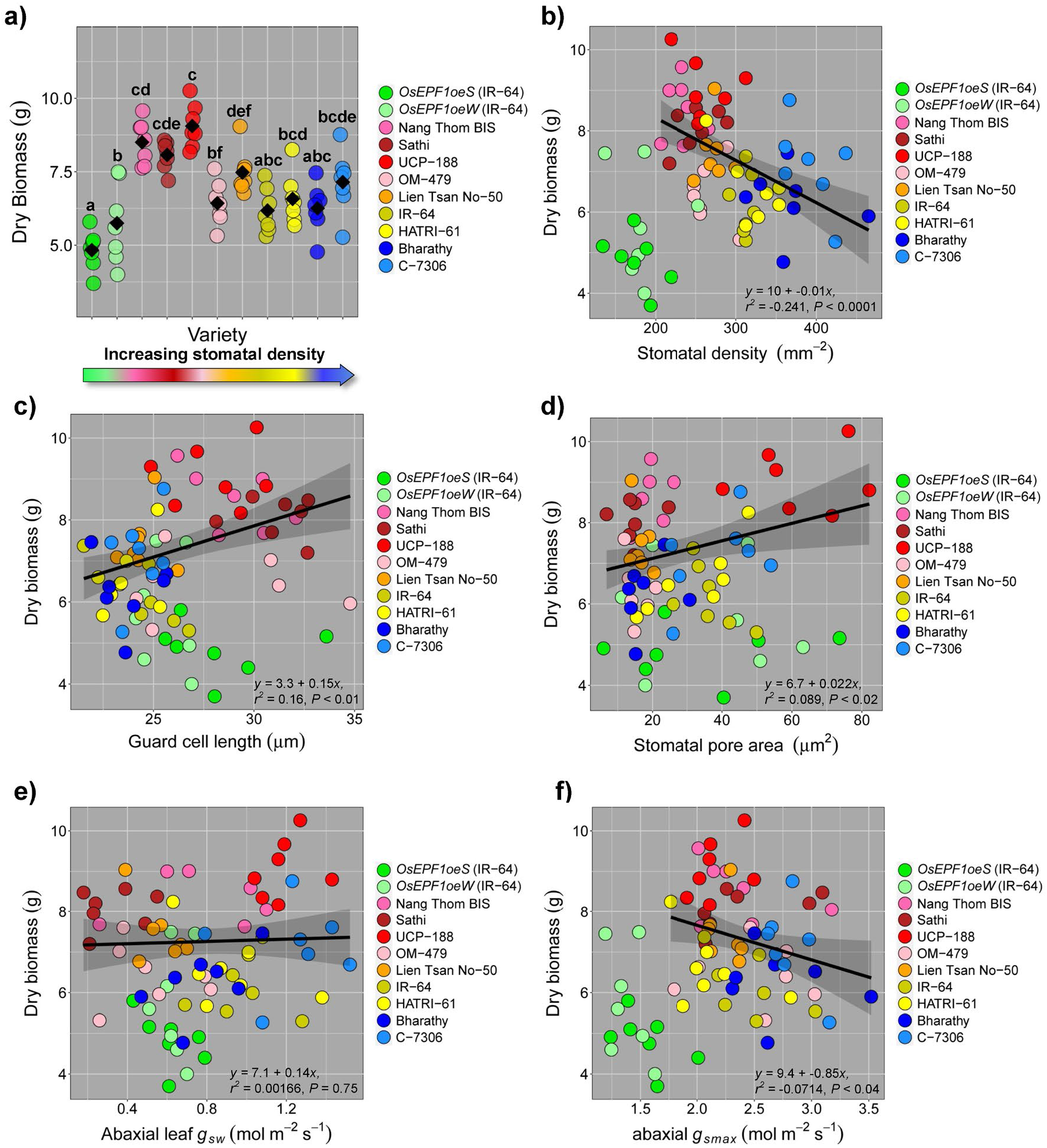
Stomatal size, density, pore area and anatomical *g_smax_* correlate with whole plant biomass during vegetative tillering. (a) Dry plant biomass. (b-f) Regression analysis conducted between biomass and (b) stomatal density, (c) stomatal size (guard cell length), (d) stomatal pore area, (e) leaf stomatal conductance (*g_sw_*), and (f) *g_smax_*. Different letters indicate a significant difference between the means (One-way ANOVA, Tukey HSD test, *P* < 0.05). Black diamonds represent means. Regression analysis and trend lines are based on linear models. *OsEPF1oe* plants are excluded from regression analyses. (a-f) *n* = 7 plants.

### The association of stomatal traits on rice resilience to drought stress

To understand how our nine selected varieties might compare to the *OsEPF1oe* plants when exposed to abiotic stress, we first imposed a drought for five days from 30 DPG (**Fig. 3**). Chlorophyll fluorescence measurements were taken to assess the efficiency of Photosystem II (ΦPSII) using a drop in the ΦPSII value to indicate plant stress (Caine *et al*., 2019).

**Fig. 3.**
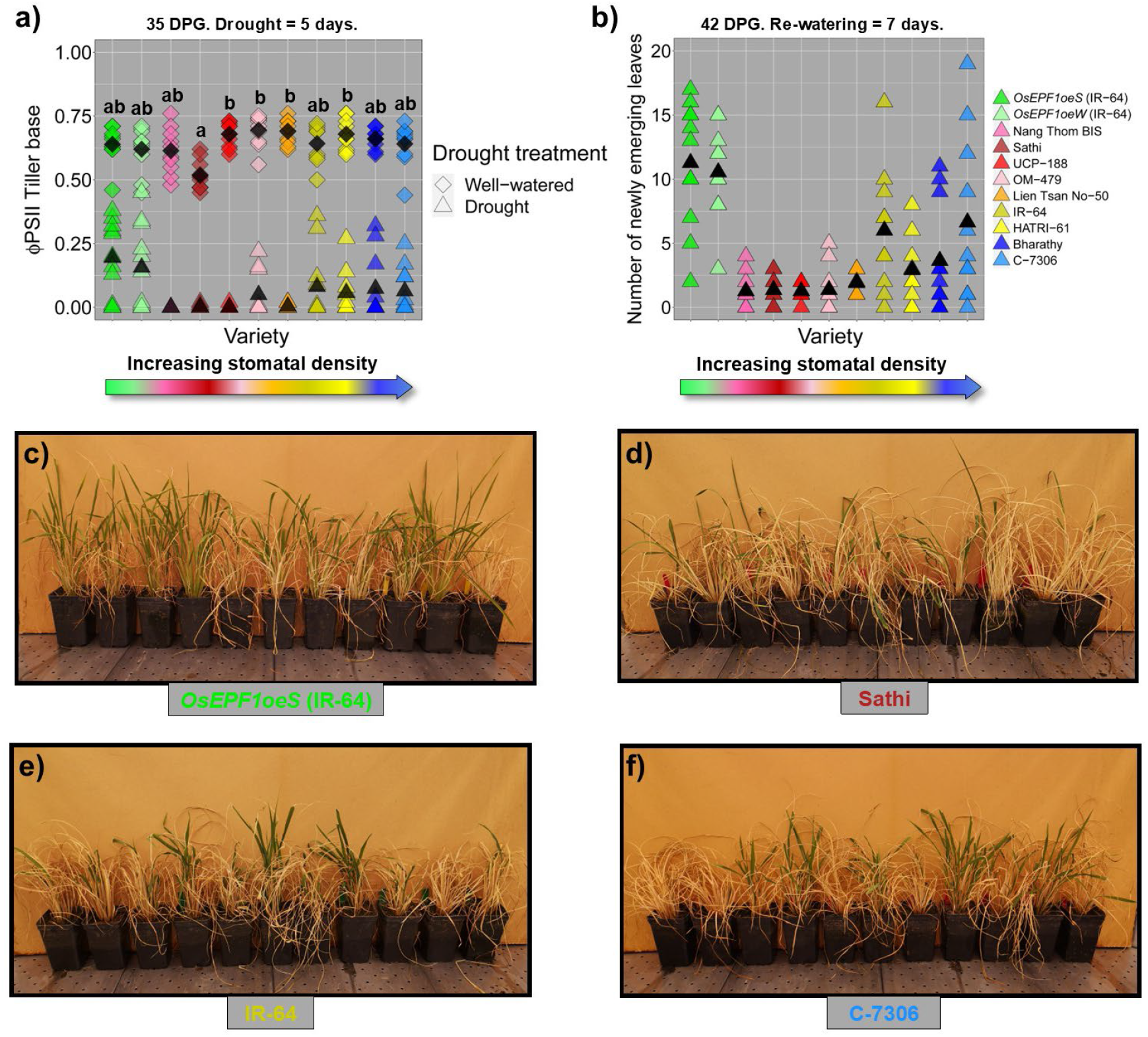
Varieties with higher stomatal density (SD) and smaller stomatal size (SS) respond better to drought than those with lower SD and large SS. (a) Drought responses assessed at the tiller base using Φ PSII to measure plant health. (b) Number of new regenerative leaves one week after re-watering. (c-f) Examples of varieties with (c) very low SD (*OsEPF1oeS*), (d) low SD and large SS (Sathi), (e) medium SD and small SS (IR-64) and, (f) high SD and small SS (C-7306) recovering from drought at 42 days, one week after re-watering. Different letters indicate a significant difference between the means (One-way ANOVA, Tukey HSD test, *P* < 0.05). Black diamonds represent means for well-watered plants, and triangles represent means for droughted plants. (a-b) *n* = 10-11 plants.

Under well-watered conditions, the *OsEPF1oe* lines and most of the selected varieties shared similar ΦPSII values at the tiller base, with only Sathi displaying slightly (but significantly) lower values than 4 of the other varieties. Following five days of severe drought treatment the ΦPSII values of all traditionally-bred varieties and *OsEPF1oe* lines were dramatically lower than in well-watered controls. Varieties previously identified as having low SD (mostly with large stomata: Nang Thom BIS, Sathi, UCP-188 and OM-479) did not perform well, with most exhibiting ΦPSII values of 0 at the end of the drought (36 out of 44 plants), suggesting that the tillers of these plants were severely stressed (**Fig. 3a**). Plants previously identified to have medium to high SDs (typically with smaller SS: IR-64, HATRI-61, Bharathy and C-7306) performed better, with just over half of plants recording ΦPSII values of 0 (25 out of 44). Although there was variability between individual plants, the *OsEPF1oe* lines fared best and maintained the highest mean ΦPSII values (only 4 out of 22 plants displayed a ΦPSII value of 0).

All plants were then resupplied with water and new leaf growth was assessed after 7 days (**Fig. 3b-f**). This revealed that the 35 DAG ΦPSII measurements were a good indicator of plants drought resilience. Plants with low densities of large stomata recovered most slowly and had the fewest new leaves (averaging 1-2 new leaves). Medium and high SD varieties recovered better (averaging 3-7 new leaves) and the transgenic *OsEPF1oe* plants had the highest drought tolerance (averaging 10-11 new leaves per plant).

### OsEPF1oe lines have enhanced salt tolerance

To understand if stomatal traits can influence how plants perform under saline conditions, we compared the performance of the *OsEPF1oe* to IR-64 control plants grown in 20 mM NaCl.

Growth under saline conditions resulted in an increase in mean SD (19-21 DPG), with salt-treated *OsEPF1oeW* seedlings having significantly more stomata mm^-2^ than equivalent fresh-water grown plants when leaf 5 seedlings were assessed (**Fig. 4a**). SS was unaffected by salt treatment, although*OsEPF1oeS* plants had the smallest SS when grown in fresh water or 20 mM NaCl (**Fig. 4b**) (this is more similar to SS trends previously reported for seedling leaves by Caine et al (2019)., than that reported for leaf 8 of tillering plants in **Fig. 1b**). Gas exchange analysis showed that *OsEPF1oeS* (but not *OsEPF1oeW*) had significantly reduced *A* and *g_sw_* relative to IR-64 when grown in fresh-water (**Fig. 4c** and **d**). Under saline conditions, IR-64 *g_sw_* was greatly reduced, and the *A* and *g_sw_* rates were no longer significantly higher than either *OsEPF1oe* line (**Fig. 4c** and **d**). However, the (already low) *g_sw_* of *OsEPF1oe* plants remained relatively similar between fresh water controls and salt-treated equivalents. The reduced *A* and *g_sw_* of salt-treated IR-64 plants resulted in an increased intrinsic water-use efficiency (iWUE), whereas for *OsEPF1oe* plants, iWUE did not significantly increase under saline conditions (**Fig. 4e**).

**Fig. 4.**
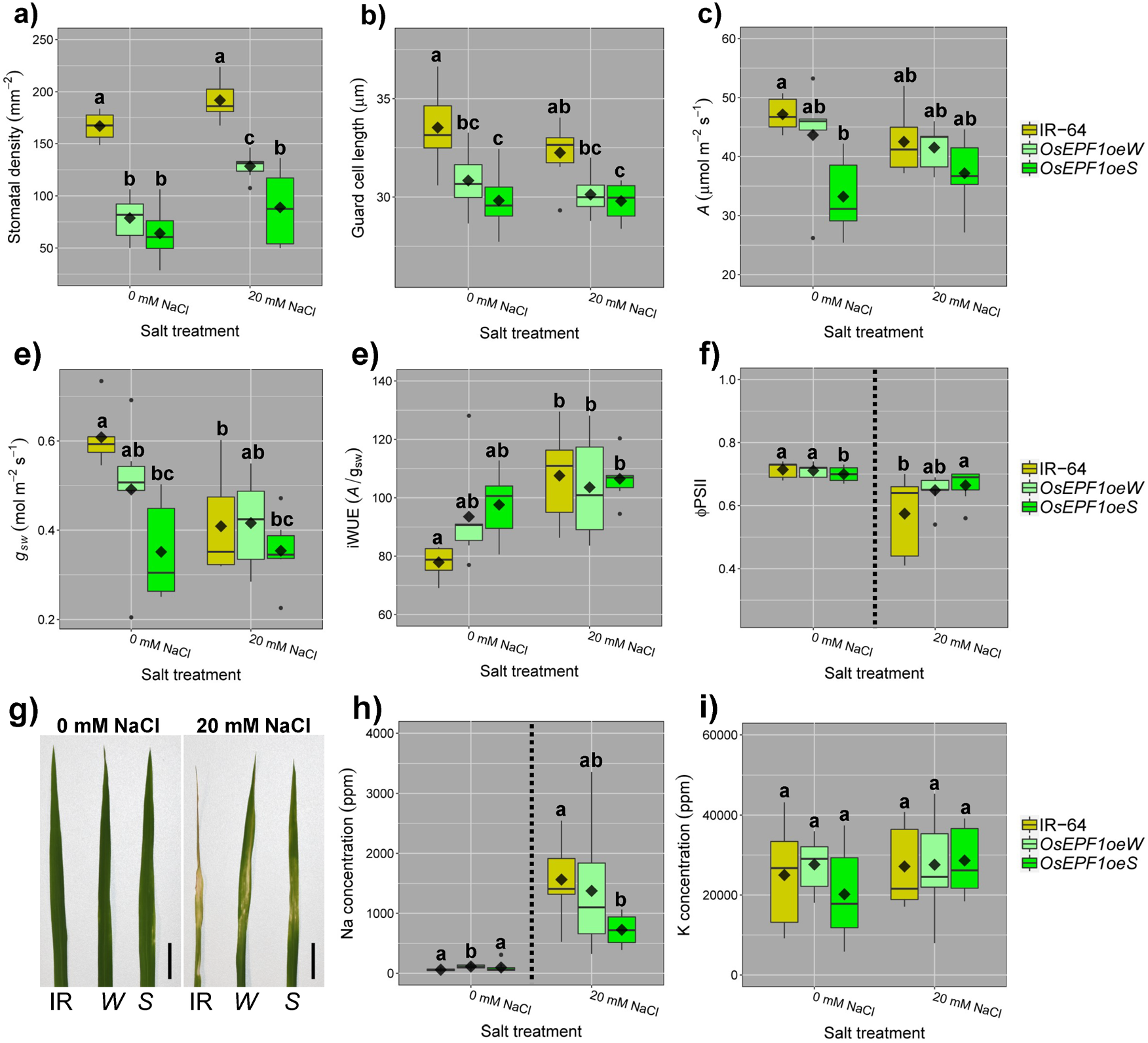
*OsEPF1oe* plants with reduced stomatal density (SD) display increased salinity tolerance during seedling and tillering stages. (a) 19-21 DPG Leaf 5 SD and (b) Stomatal size (guard cell length) of fresh water and salt grown IR-64 and *OsEPF1oe* plants. (c-f) Fresh water and salt grown rice plant gas exchange measurements of (c) Assimilation (*A*), (d) stomatal conductance (*g_sw_*) and (e) intrinsic water-use efficiency (iWUE). (f) Φ PSII leaf measurements of apical leaves at 28 DPG with (g) representative leaf images from fresh water and salinity treated plants (bars, 2 cm). (h) Sodium (Na), and (i) Potassium (K) concentrations in auxiliary leaves of 35 DPG tillering plants. Whiskers indicate the ranges of the minimum and maximum values and different letters indicate a significant difference between the means (Two-way ANOVA, Tukey HSD test, *P* < 0.05). For (f) and (h), 2 separate Kruskal-Wallis one-way ANOVAs were performed due to unequal variances (*P* < 0.05). Black diamonds represent means and black dots are outliers. *n* = 6-7 plants.

Plants were left to develop further and at 28 DPG the continuing impact of salt uptake was investigated using ΦPSII values as a proxy for plant health (**Fig. 4f**). Under normal fresh water conditions the ΦPSII of *OsEPF1oeS* (but not *OsEPF1oeW*) leaves were was significantly lower than IR-64. Salinity treatment had a more severe impact on IR-64 than the *OsEPF1oe* lines and salt-treated *OsEPF1oeS* had higher ΦPSII values than IR-64, with visibly healthier leaves (**Fig. 4f-g**). At 35 DPG the concentration of accumulated salt in auxiliary leaves was measured. The plants with the lowest SD accumulated significantly less salt in their leaves, with *OsEPF1oeS* salt-grown plants having c. 50% lower Na than IR-64 equivalents (**Fig. 4h**). In comparison, we did not detect any differences in K levels across genotypes or treatments (**Fig. 4i**).

Next, we investigated how the nine selected traditionally-bred varieties and two transgenic lines performed under exposure to 50 mM NaCl from 16 DPG onward. We measured ΦPSII every 3 to 4 days for the next 67 days (**Fig. 5**).

**Fig. 5.**
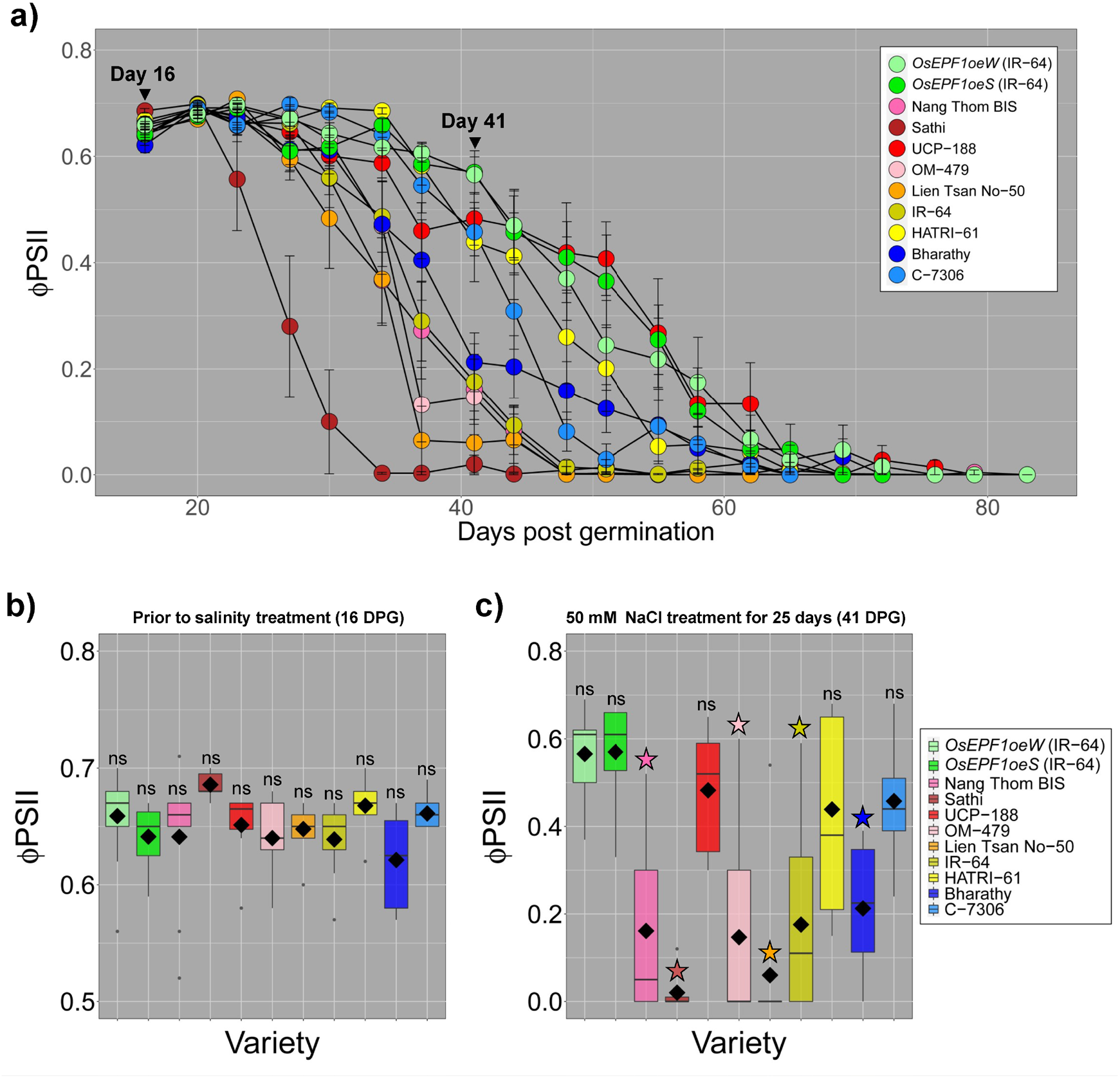
*OsEPF1oe* plants maintain higher leaf ΦPSII values for equal or longer than all traditionally-bred varieties. (a) Apical leaf ΦPSII measurements of the nine selected varieties and two *OsEPF1oe* lines grown in 50 mM NaCl solution for 67 days. Vertical lines above and below individual points show one standard error (b) There was no statistically significant differences between *OsEPF1oeW* ΦPSII and the other traditionally-bred varieties or the *OsEPF1oeS* line prior to plants beginning the salt treatment at day 16 (denoted ns). (c) ΦPSII values on the apical leaf at 41 DPG, 25 days after the commencement of the salt treatment. For (b) and (c), whiskers indicate the ranges of the minimum and maximum values and stars indicate a significant difference from salt tolerant *OsEPF1oeW*(One-way ANOVA, Tukey HSD test, *P* < 0.05). Black diamonds represent means. *n* = 7-9 plants.

Prior to the start of salt treatment (16 DPG), no significant differences between control *OsEPF1oeW* leaf ΦPSII values and any of the other varieties tested (**Fig. 5a, b**). After 25 days of salt treatment (41 DPG), six out of nine varieties (including IR-64) showed significant reductions in ΦPSII (**Fig. 5a, c**). Post 41 DPG, *OsEPF1oe* plant leaves continued to show a slower decline in health than the majority of varieties, with the exceptions being UCP-188. Within the selected varieties, neither SS nor SD appeared to be associated with the degree of salinity tolerance.

### Stomatal responses to raising temperature and vapour pressure deficit

At higher temperatures the amount of water the air can hold increases and this often leads to rising VPD bringing about stomatal closure. We next investigated how increases in VPD impacted stomatal dynamics by undertaking gas exchange experiments that either rapidly increased VPD by increasing temperature (**Fig. 6** and **Fig. S3**), or by exposing a leaf to a constantly high VPD (**Fig. S3**). High temperature and VPD conditions were imposed inside an infrared gas analyser (IRGA) leaf chamber and gas exchange simultaneously measured. For our rapid VPD experiments, assays were conducted over a 1 hour duration, with a 10 °C increase in temperature inside the chamber (from 32 °C to 42 °C), beginning 12 minutes into the experiment. The leaf VPD inside the chamber increased from c. 2.6 kPa to 3.7 - 4.7 kPa (**Fig. 6b, Fig. S3**). Variation in leaf VPD and chamber relative humidity (RH) at higher temperatures (≥ 39.5 °C) was due to reductions in water flow caused by increased stomatal closure and by inability of the IRGA to maintain RH.

**Fig. 6.**
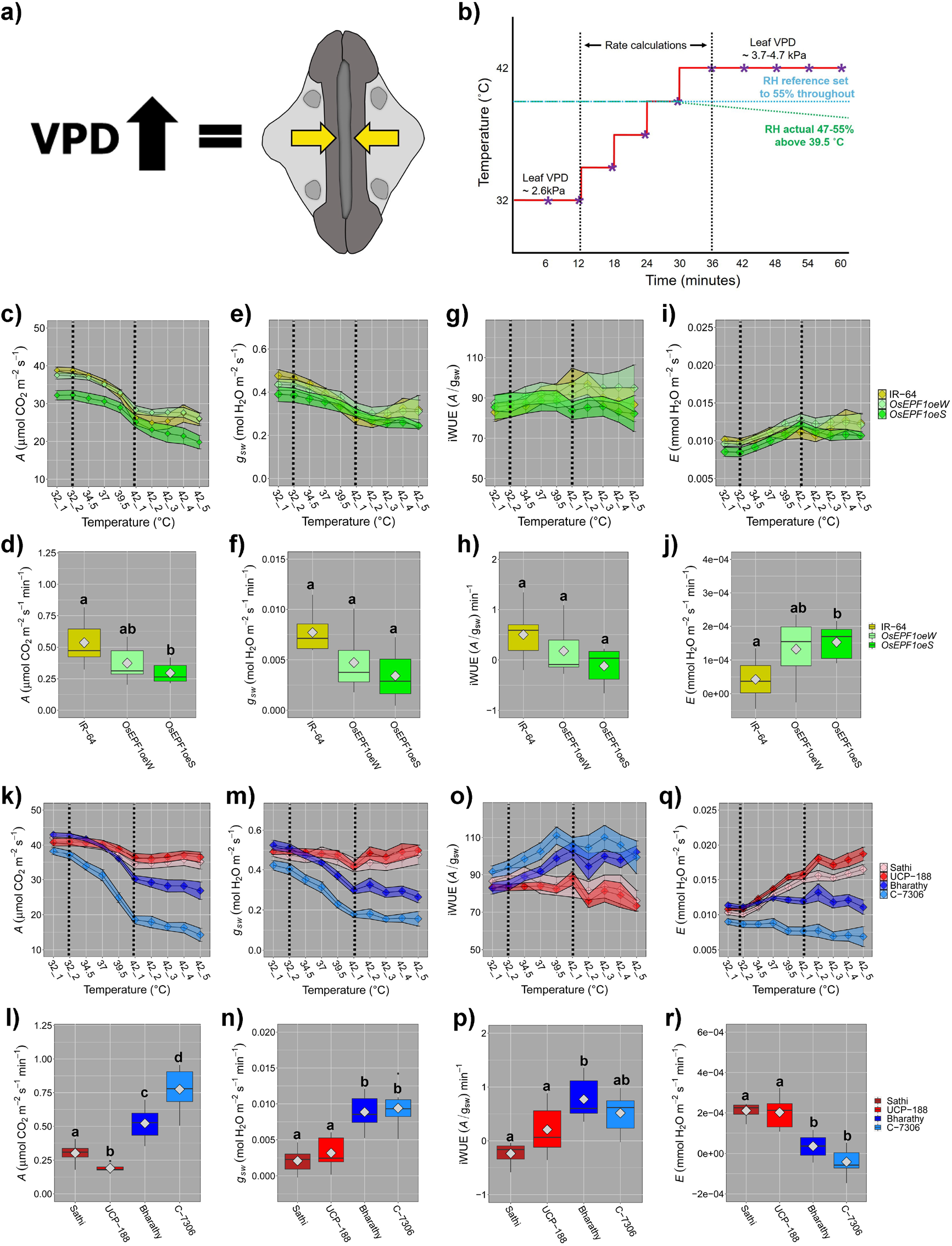
Dynamic stomatal responses to rising Vapour Pressure Deficit (VPD) lead to large alterations in plant photosynthesis and water-use efficiency. (a) When VPD rises, usually driven by increasing temperature, stomata close. (b) Program performed in LI-COR 6800 gas analysers used to study stomatal responses to temperature and leaf VPD. Around 39.5 °C and above, many varieties were unable to maintain relative humidity at 55% due to additive RH reaching its maximum possible set point (see also Fig. S3). Purple stars indicate points data was recorded. Parallel vertical dotted lines throughout indicate period used to calculate rate changes. (c-j) Comparison of IR-64 plants and *OsEPF1oeW* and *S* plants using leaf 5 to investigate responses to raising temperature and VPD. (c) Assimilation (*A*) responses over a one hour duration and (d) rate of change per minute during the 24-minute incline period. (e-f) Equivalent stomatal conductance (*g_sw_*) responses and rates of change over 24-minute period. (g-h) Corresponding intrinsic water-use efficiency (iWUE) responses and rates of change and (i-j) transpiration (*E*) responses and rates of change. (k-r) Comparisons between low SD, large SS Sathi and UCP-188 varieties and high SD, small SS Bharathy and C-7306. (k) *A* responses over one hour assay and (l) rate of change per minute during the 24-minute incline period. (m-n) Equivalent stomatal conductance (*g_sw_*) responses and rates of change. (o-p) Corresponding intrinsic water-use efficiency (iWUE) responses and rates of change and (q-r) transpiration (*E*) responses and rates of change. Ribbons highlight standard error of the mean. Boxplot whiskers indicate the ranges of the minimum and maximum values and different letters indicate a significant difference between the means (One-way ANOVA, Tukey HSD test, *P* < 0.05). Grey diamonds represent means. *n* = 7-8 plants.

We first assayed IR-64 and *OsEPF1oe* plants and found that all plants showed a significant decrease in *A* as temperature and VPD increased (ANOVA, *P* < 0.0001, **Fig. 6c**), but the reduction in *A* occurred significantly more slowly for *OsEPF1oeS* than in IR-64, but not for *OsEPF1oe*W (**Fig. 6d**). Like *A, g_sw_* also reduced with increasing temperature (ANOVA, *P* < 0.001, **Fig. 6e**), however there were no significant differences in the rate of change between genotypes, despite a clear trend for a slower rate in *OsEPF1oeS* (Kruskal-Wallis one-way ANOVA, overall P = 0.056) (**Fig. 6f**). The slower decreases in *A* and (to some extent *g_sw_*) resulted in a trend toward lower iWUE for *OsEPF1oeS* during the temperature incline relative to IR-64 (**Fig. 6g and h**) (Kruskal-Wallis one-way ANOVA, overall P = 0.1052). Conversely, transpiration (*E*) responses generally increased across all genotypes with *OsEPF1oeS* plants having increased rates of *E* relative to IR-64 during the temperature incline, peaking at the end of the increase. The *E* of IR-64 peaked later and by the end of the 1 hour experiment reached a similar level to *OsEPF1oe* plants (**Fig. 6i-j**).

To investigate if a combination of SS and SD contributed to rice VPD responses, we compared two selected varieties which had low SD and large SS (Sathi and UCP-188), with two varieties which had high SD and low SS (Bharathy and C-7306) (**Fig. 6k-r** and **Fig. S3**). There were no significant differences in *A* or *g_sw_* at 32 °C between the two low SD, large SS varieties and the two high SD, small SS varieties at the beginning of the experiments, but large differences were detectable in response to rising temperature and VPD (**Fig. 6k-n**). Specifically, varieties with high SD with small SS showed significantly faster rates of reduction for both *A* and *g_sw_* in response to rising temperature and VPD in comparison to varieties with low SD and large SS (**Fig. 6l-n**). This faster response resulted in a large increase in iWUE (**Fig. 6g, o-p**). We also detected striking differences in *E* in response to rising VPD, with varieties with low SD and large SS displaying a rapid increase in *E* with increasing temperature which was in contrast to plants with high SD and small SS that showed little change (or a slight drop) in *E* (**Fig. 6q-r**). Thus, the two varieties with higher SD and smaller SS reduced *A* and *g_sw_* relatively quickly in response to an increase in VPD whereas for low SD varieties with large SS, *A* and *g_sw_* remained higher and this led to increasing *E*.

The experiments in Fig. 6 highlight that high SD, small SS rice varieties were unable to maintain high *A, gsw and E* when exposed to increasingly high temperature and VPD. It is possible that that this may have, at least in-part, been due to a transient drop in RH within the chamber during the rapid closure of small stomatal that contributed to leaves having higher VPD (represented by green dotted line **Fig. 6b,** see also **Fig. S3**). To address this, we conducted a subsequent experiment where we held plants under steady-state conditions with temperature fixed at 39 °C where chamber RH was maintained at stable at 55%, and this led to more similar leaf VPD values between varieties (**Fig. S3**). These conditions captured the maximum point where all 4 different rice varieties were able to maintain steady state conditions. This removed any additional stress on the plant leaves caused by insufficient supply of RH as observed in our Fig. 6 experiments. Like with our dynamic response Fig. 6 experiments, plants with low SD and large SS maintained a higher A and *g_sw_* than those with high SD and small SS. Reductions in *A* and *g_sw_* again had the opposite effect on iWUE, with high SD, small SS plants typically having higher iWUE at the expense of *A* and *g_sw_* (**Fig. S3**). And also like with our dynamic response experiments in Fig. 6, these changes appeared to be entwined with an increased capability of plants with low SD and large SS plants to increase *E*, whereas for plants with high SD and small SS, *E* stayed low. These combined responses led to plants with lower SD and larger SS having lower leaf temperatures based on calculated energy balance (**Fig. S3**).

## Discussion

### Stomatal size and density impact on plant gas exchange

A low SD (often associated with larger SS) and/or small SS (often associated with high SD) have frequently been correlated with improvements in water-use and/or drought tolerance (Hepworth *et al*., 2015; McAusland *et al*., 2016; Dittberner *et al*., 2018; Kardiman & Ræbild, 2018; Caine *et al*., 2019; Mohammed *et al*., 2019). In this study, we took a stomatal-focused approach to investigate how differences in SS and SD can affect rice performance under several separate abiotic stresses. Anatomical screening of a rice population identified significant variation in both traits across the population. We then compared the abiotic stress responses of selected varieties with a range of differences in SS and SD and *OsEPF1oe* plants which have an unusual combination of low SD and small SS (on their seedling leaves). None of the varieties examined had an SD as low as *OsEPF1oe* lines, suggesting that finding traditionally-bred rice varieties with SDs and *g_smax_* values equivalent to *OsEPF1oe* is perhaps unlikely.

Previous research has shown that high SD (often accompanied by small SS) can lead to a higher maximum anatomical *g_smax_* (Franks & Beerling, 2009), and this leads to a higher operating *g_sw_* and greater responsivity of stomatal apertures to environmental changes (Franks *et al*., 2012; Bertolino *et al*., 2019). Our results only partially support these findings. We observed the expected correlations between SD and both *g_smax_* and *gsw;* with the two varieties with highest SD and small SS (Bharathy and C-7306) having the highest average *g_smax_*. We did not, however, find a correlation between operating *g_sw_* and *g_smax_* under well-watered conditions underlining that factors other than SD and SS, including stomatal openness, also impact on operating *g_sw_*. For example, the UCP-188 variety with low SD and large SS, despite having a low calculated *g_smax_*, had the equal highest operating *g_sw_*, and *OsEPF1oe* lines with only 50-72% of the *g_smax_* of the traditionally-bred varieties, had an operating *g_sw_* similar to other low SD varieties (except UCP-188). Indeed, plotting stomatal pore area measurements against *g_sw_* indicated that *OsEPF1oe* plants can counteract their reduced SD by increasing stomatal pore areas (**Fig. 1i**).

Our observed correlations between plant biomass and SD, and plant biomass and SS suggest that stomatal characteristics at the tillering stage are reasonable predictors of plant size (**Fig. 2**). Plants with low SDs and large SS typically had higher biomasses (with the exception of *OsEPF1oe*), and plants with a high SD and small SS typically had lower biomasses. Importantly, differences in biomass have the potential to exacerbate differences in water-use, as plants with a larger surface area often require more water (Feldman *et al*., 2018) and may also close their stomata more slowly than smaller plants (Drake *et al*., 2013; McAusland *et al*., 2016; Lawson & Vialet-Chabrand, 2019).

### Stomatal associations with drought tolerance

Our drought stress experiments were designed to investigate if larger plants with large SS displayed slower stomatal responsiveness to water stress (**Fig. 3**). We found that traditionally-bred varieties, with high or medium SD with small SS, maintained higher ΦPSII values for longer during drought than varieties with low SD and large SS, indicative of better plant health. However, this was not the case for the *OsEPF1oe* plants. Both lines of these very low SD plants had lower biomass, and these combinations of traits resulted in *OsEPF1oe* plants maintaining the highest ΦPSII at the end of the drought. This was despite *OsEPF1oe* plants showing similar operating *g_sw_* levels as other varieties with low SD, large SS when assessed under well-watered conditions (**Fig. 1e**). These results suggest that the ideal combination of traits for growth of rice under drought conditions, would be small plant size together with either a very low SD (leading to a very low *g_smax_*), or a higher SD combined with a small SS (leading to a high *g_smax_* and greater stomatal responsivity).

### Stomatal contributions to salinity tolerance

Modelling of root water and solute uptake has suggested that passive transport and uptake of water and solutes into plants may be negligible (Foster & Miklavcic, 2017). However, recent research in rice has suggested that reduced SD and gsw, caused by increased activity of a histone deacetylase, improved both drought and salinity tolerance (Zhao *et al*., 2021). In our study, we tested this potential relationship between SD and salt tolerance and found that *OsEPF1oe* plants with reduced SD showed improved salinity treatment (**Fig. 4**). We found that these plants with maintained higher leaf ΦPSII values when grown in salt water, and also accumulated less than half the amount of Na^+^ in leaves after approximately 5 weeks of growth (**Fig. 4**). Salt toxicity often leads to deficiencies in other elements such as K^+^ (Wang *et al*., 2013) but we did not detect this in *OsEPF1oe* plants or controls. We compared the performance of traditionally-bred varieties against *OsEPF1oe* and found that, as in the drought experiments, *OsEPF1oe* plants performed well, maintaining leaf ΦPSII the equal longest of all varieties surveyed. In contrast, other varieties with low SD and large SS performed the least well (with the exception of UCP-188) (**Fig. 5a, c**).

### Stomatal responsiveness to rising temperature and VPD

Rising temperatures leading to increased VPD has the potential to shut stomata at a time when plants might otherwise utilise transpiration-driven evaporative cooling to maintain a high photosynthetic output (Urban *et al*., 2017; Yuan *et al*., 2019; Grossiord *et al*., 2020). Comparisons between IR-64 and transgenic plants revealed that *OsEPF1oeS* seedlings had slower rates of photosynthetic decline at higher temperature and VPD, and this was linked with a trend towards a decreased rate of *g_sw_* change per minute (**Fig. 6**). These slower reductions in *A* and operating *g_sw_* were coupled with increased rates of *E*, suggesting that *OsEPF1oeS* leaves with very low SD lost more water in comparison to IR-64 as VPD stress increased. This slower response suggests that having vastly reduced stomata may be detrimental when temperature and VPD are constantly fluctuating, but this requires further study to confirm.

Small SS in nature is often associated with faster stomatal responsiveness, which in-turn can result in higher water-use efficiency under fluctuating conditions (McAusland *et al*., 2016; Lawson & Vialet-Chabrand, 2018; Inoue *et al*., 2021). However, if plants with small SS were to close stomata rapidly on sensing high VPD, this could lead to detrimental reductions in *A* and evaporative cooling. We therefore tested whether varieties with small SS had faster or slower stomatal VPD responses by comparing two varieties with the highest SD and small SS with two varieties which had lowest SD and large SS (**Fig. 6** and **Fig. S3**). We found that, unlike *OsEPF1oe* lines, the varieties with small SS (and higher SD) reduced their *g_sw_* much more rapidly than those with large SS. Indeed, the extremely efficient closure of small SS (and high SD) reduced *g_sw_* so effectively that the IRGA equipment measuring the plants was unable to maintain RH levels within the leaf chamber, whereas this was not apparent for plants with larger SS (and low SD) (**Fig. S3c**). This rapid stomatal closure in plants with small SS resulted in improved iWUE at higher VPD levels, but this was at the expense of *A*. In contrast, the varieties with larger SS maintained higher rates of *A* and *g_sw_* fairly steadily with increasing temperature and VPD, but their level of *E* increased markedly. These results illustrate how differing stomatal configurations are associated with differing responses; small SS with high SD can react more quickly to reduce *g_sw_* whereas plants with larger SS and low SD are usually less responsive in controlling their water loss via stomata at high temperature and VPD. These differing stomatal strategies most probably allow plants to inhabit and thrive under different environments. For example, the fast responses of small stomata (potentially with high SD) would be expected to be beneficial in water restricted conditions, but higher levels of *E* associated with plants with larger SS (potentially with lower SD) could have a greater effect on the microenvironment. This may be a positive trait when growing in high temperature or VPD environments if sufficient water is available to power an enhanced transpiration stream.

### Effects on Stomatal Development

Our results support previous observations (Zhang et al., 2019) showing that rice SD and SS are, in general, negatively correlated (**Fig. 1 and Fig. S1**). During leaf development, epidermal cell division is usually synchronised with cell expansion to achieve a specific SS and SD on a given leaf. This relationship is perturbed in *OsEPF1oe* seedlings, where plants with extremely low SD, typically have smaller SS at the seedling stage (Caine *et al*., 2019). However, as we show here, by leaf 8 stage when plants are rapidly tillering, this is no longer the case, and *OsEPF1oe* plants had comparatively larger SS (than IR-64) in their more mature leaves. SD is also under developmental control, and in our experiments the leaves of mature plants had considerably more stomata than seedling leaves. Thus, the potency or nature of factors driving SS and/or SD regulation must change as a plant develops. Recently GWAS analysis has identified genetic traits associated with both SS and SD which could be useful in breeding for these traits (Chen *et al*., 2020).

## Conclusion

Previously we have shown that *OsEPF1oe* plants with extremely low SD, have improved drought tolerance and survivability, even at high temperatures. Here, we show that these plants are also less susceptible to salinity toxicity, most probably because they accumulate salt at much lower levels. By screening a range of traditionally-bred rice varieties, we show that varieties with low SD and large SS typically have lower operating *g_sw_* when sufficient water is available, but this is often associated with increased biomass that could promote increased drought susceptibility. *OsEPF1oe* plants were physically smaller and yet had the lowest SD and *g_smax_* of all varieties studied, traits which together appeared to positively impact on both drought and salinity tolerance. Based on our screen, it seems unlikely that similar conformations of stomata with correspondingly small plant size will be found in traditionally-bred rice varieties. This means that stomatal-based stress tolerance may best be obtained via genetic manipulation of stomatal development rather than via conventional breeding practices.

Assessment of VPD responses suggest that strong over-expression of *EPF1* diminishes stomatal closure responses relative to IR-64 leading to reduced short-term iWUE. This slower stomatal responsiveness leads to increased *E*. Conversely, traditionally-bred plants with high SD and small SS greatly reduce *g_sw_* at high VPD, preventing *E* increasing, and this may be beneficial for short-term iWUE, but detrimental to long-term plant productivity and cooling. Whilst *OsEPF1oe* plants showed enhanced tolerance to several abiotic stresses, taken together our results highlight that there is unlikely to be an ideal SS and SD for all future climatic conditions, and thought must be given as to which conformation(s) of stomata will best suit a given growth environment.

## Acknowledgments

We thank Dr Jose De Vega, Earlham Institute, for providing the rice germplasm. We are grateful for funding from BBSRC Newton Fund (BB/N013646/1) and a Leverhulme Trust Senior Research Fellowship (SRF\R1\21000149) to JEG, a University of Sheffield QR GCRF Fellowship (Research England institutional allocation) to RSC, and a BBSRC DTP studentship to ELH (BB/M011151/1). HLC was supported by the UK Research and Innovation (UKRI) Future Leaders Fellowship scheme [MR/T01993X/1]

## Author contributions

R.S.C., N.T.L., J.E.G. designed the study. R.S.C., E.L.H., J.M.S., P.M.F., and S.F. undertook the experiments. N.T.L., N.T.P., H.L.C. and J.E.G. contributed materials and advice. R.S.C., H.L.C. and J.E.G. wrote the paper with comments from E.L.H, J.M.S., P.M.F., S.F., N.T.L. and N.T.P. All authors read, commented on and approved the final version of the manuscript.

---

**Fig. S1.** Rice leaf 5 stomatal size and density screen of 72 traditionally-bred rice varieties and 2 transgenic varieties.

**Table. S1.** List of 72 rice varieties and corresponding stomatal size and densities.

**Fig. S2.** Stomatal size and density relationships to stomatal conductance.

**Fig. S3.** Rapid and steady-state VPD responses of rice with differing stomatal size and density.

## Notes

### Competing Interest Statement

The authors have declared no competing interest.

