## Supplemental information for "The Influences of Stomatal Size and Density on Rice Drought, Salinity and VPD Resilience"

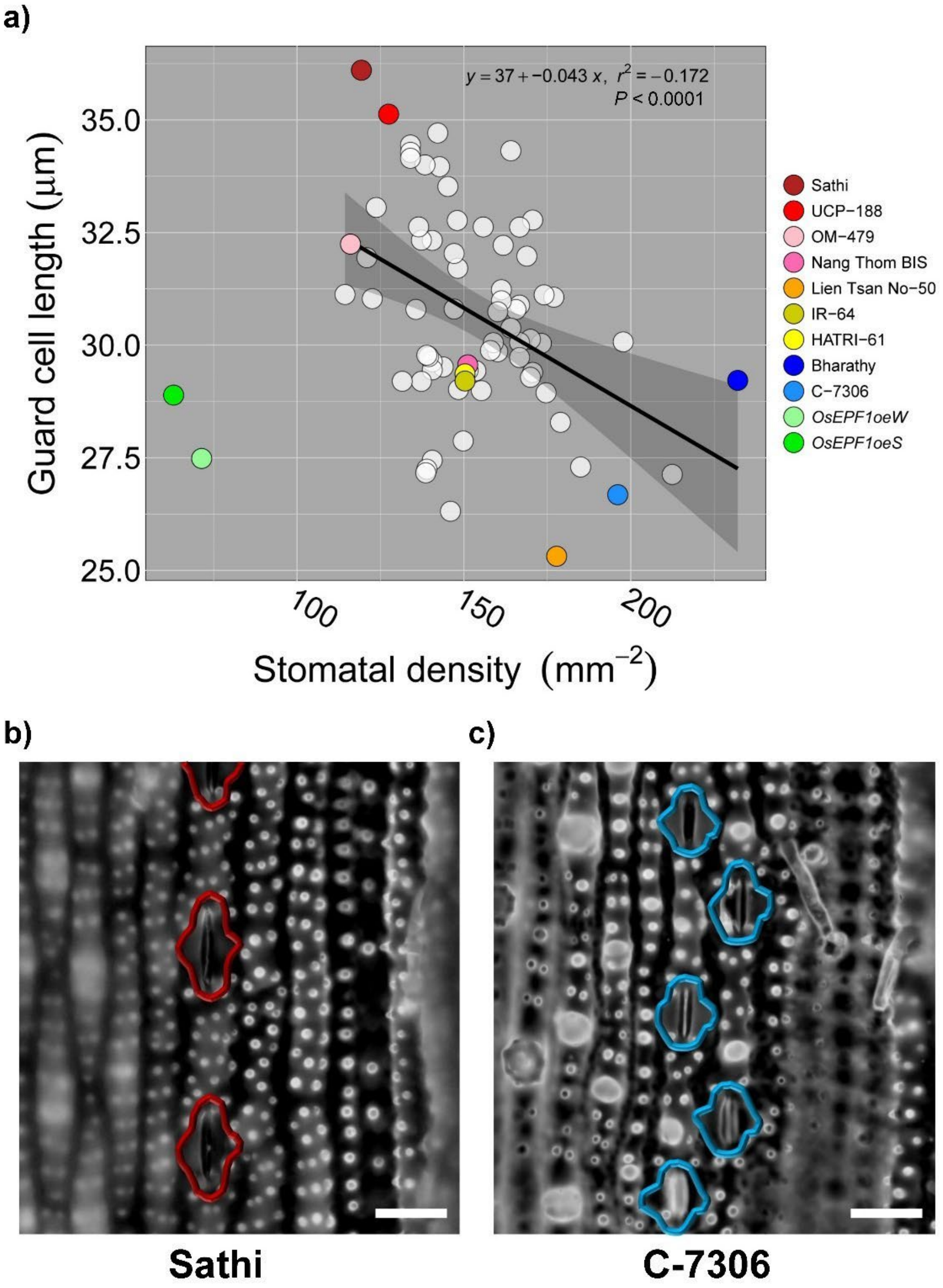


**Fig. S1.** Screening for stomatal size and density variation amongst 72 traditionally-bred bred rice varieties and 2 transgenic *OsEPF1oe* lines on leaf 5. (a) Stomatal size and density plot of assessed varieties and transgenics. 9 varieties and 2 transgenic lines assessed in the manuscript for gas exchange and abiotic stress performance are highlighted by different coloured circles. White circles denote other screened varieties. See also Table S.1 for a full list of varieties. (b-c) Representative epidermal images of stomatal size and density differences (b) low stomatal density, large size Sathi and (c) high stomatal density, small size C-7306 (bar, 25 μm).

**Table S1.** Stomatal size and density measurmentes of 72 varieties and 2 transgenic lines assessed during screen. *n = 1-7*

| Variety | Stomatal density ($mm^{-2}$) | Guard cell length (µm) | Replicate number | Variety | Stomatal density ($mm^{-2}$) | Guard cell length (µm) | Replicate number |
|---|---|---|---|---|---|---|---|
| Bharathy | 232.0 | 29.22 | 4 | OM-5629 | 149.8 | 27.87 | 6 |
| Tep Hanh | 212.4 | 27.13 | 1 | Doc Phung Lun | 148.4 | 29.02 | 5 |
| Nang Du | 197.7 | 30.07 | 2 | HATRI-20 | 148.1 | 31.71 | 6 |
| C-7306 | 196.1 | 26.68 | 2 | HATRI-192 | 148.0 | 32.77 | 7 |
| Cadung Gocong | 185.0 | 27.30 | 5 | Kharai Ganga | 147.1 | 32.03 | 1 |
| IARI-5823 | 178.9 | 28.29 | 4 | Mot Bui Do | 147.1 | 30.80 | 1 |
| Lien Tsan No-50 | 177.8 | 25.31 | 5 | OM-442 | 146.0 | 26.31 | 3 |
| Gogo Putih | 176.9 | 31.06 | 7 | Jao Hom Nin | 145.2 | 33.52 | 7 |
| Nang Nhen Thom | 174.6 | 28.94 | 7 | CTG-1516 | 143.8 | 29.52 | 3 |
| DNJ-155 | 174.0 | 31.12 | 4 | OM-447 | 142.7 | 33.96 | 3 |
| San Tan Thou | 173.2 | 30.04 | 6 | Nang Thom | 142.2 | 34.71 | 2 |
| DNJ-11 | 170.6 | 32.78 | 5 | Nang Rum Trang | 140.5 | 32.32 | 1 |
| ARC-10362 | 170.6 | 29.39 | 5 | OM-457 | 140.5 | 29.63 | 1 |
| Hea Doh-4 | 169.9 | 29.29 | 4 | OM-453 | 140.5 | 29.45 | 1 |
| OM-6976 | 169.9 | 30.13 | 2 | HATRI-35 | 140.5 | 27.44 | 6 |
| HATRI-31 | 168.8 | 31.98 | 3 | OM-472 | 139.4 | 29.74 | 3 |
| Nang Quot | 166.7 | 30.06 | 5 | OM-456 | 138.9 | 29.77 | 2 |
| Dinlaga | 166.7 | 30.89 | 1 | Suhasini | 138.9 | 27.25 | 2 |
| OM-9921 | 166.7 | 29.72 | 1 | OM-4900 | 138.6 | 27.17 | 5 |
| Langmanbi | 166.7 | 32.62 | 2 | Jhona-349 | 138.3 | 34.00 | 6 |
| Sarjoo-50 | 165.6 | 30.79 | 6 | Kalubala Vee | 137.3 | 32.33 | 4 |
| Khao Pahk Maw | 164.1 | 34.32 | 5 | Kakuya | 137.3 | 29.20 | 1 |
| Perrum Karruppan | 164.1 | 30.38 | 5 | Shiratama | 136.4 | 32.62 | 4 |
| Malagpit Pirurutong | 161.8 | 32.22 | 6 | 3263 | 135.6 | 30.79 | 2 |
| Doc Phung | 161.2 | 31.23 | 3 | Nang Bang Bentre | 134.0 | 34.44 | 1 |
| OM-455 | 161.2 | 30.99 | 3 | DV-86 | 134.0 | 34.28 | 1 |
| OM-476 | 160.1 | 29.85 | 1 | OM-471 | 134.0 | 34.15 | 3 |
| HATRI-608 | 160.1 | 30.74 | 2 | Tunsart Thai-Lan | 131.5 | 29.20 | 4 |
| HATRI-50 | 158.8 | 30.05 | 5 | UCP-188 | 127.5 | 35.13 | 3 |
| DHARIA | 158.0 | 29.88 | 6 | Nipponbare | 123.7 | 33.06 | 7 |
| HATRI-603 | 155.8 | 32.62 | 3 | LM1 Tanh Binh | 122.5 | 31.03 | 6 |
| Yeh Hua Chan | 155.2 | 28.99 | 4 | OM-446 | 120.9 | 31.94 | 1 |
| Pilang Baybay | 153.6 | 29.43 | 1 | Sathi | 119.3 | 36.10 | 6 |
| OM-2517 | 151.3 | 29.44 | 7 | OM-479 | 116.0 | 32.24 | 2 |
| Nang Thom Bis | 151.1 | 29.56 | 4 | OM-11735 | 114.4 | 31.12 | 1 |
| HATRI-61 | 150.3 | 29.36 | 6 | OsEPF1oeW (IR-64) | 71.4 | 27.48 | 7 |
| IR-64 | 150.3 | 29.20 | 6 | OsEPF1oeS (IR-64) | 63.0 | 28.89 | 7 |

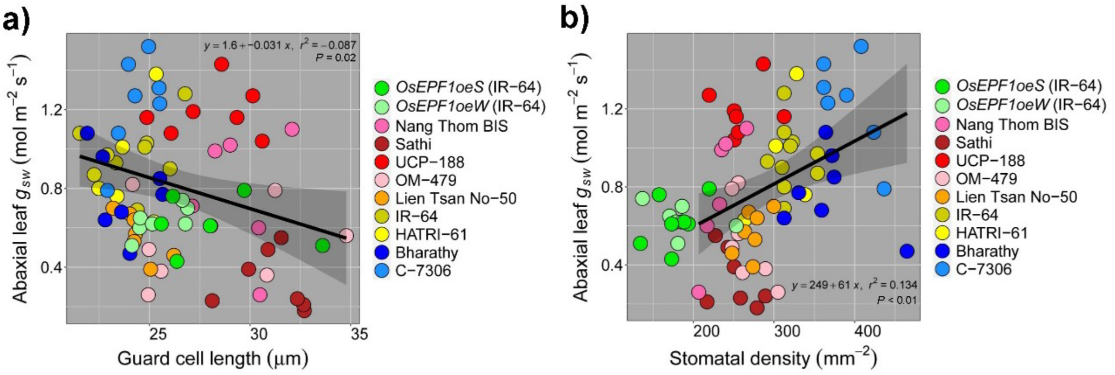


**Fig. S2.** Gas exchange relationships linked to stomatal size and density (a) Regression analyses conducted between (a) Abaxial leaf $g_{sw}$ and stomatal size (guard cell length) and (b) abaxial $g_{sw}$ and stomatal density. Regression analysis and trend lines are based on linear models. *OsEPF1oe* plants are excluded from regression analyses. (a-b) $n$ = 7 plants.

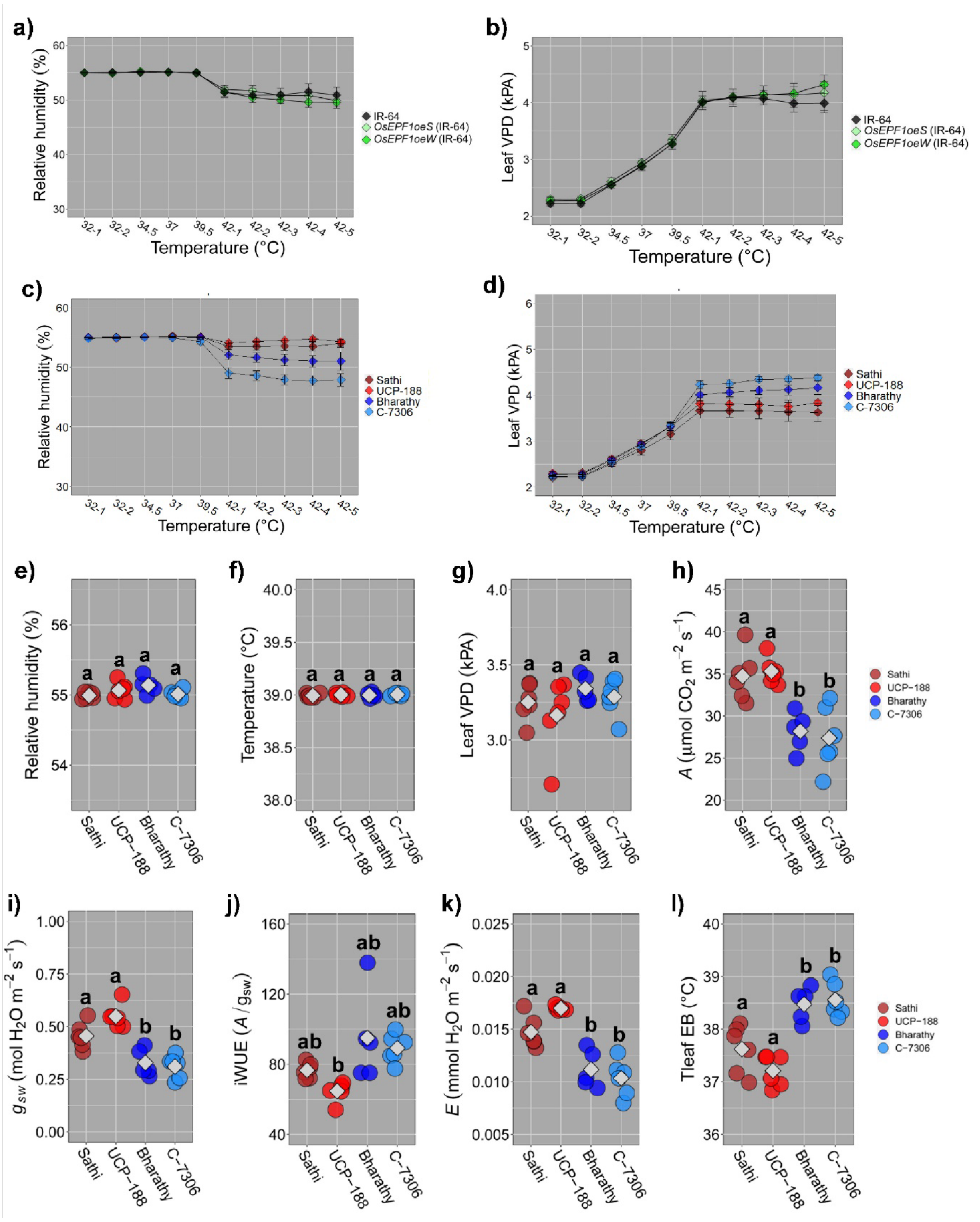


**Fig. S3.** High VPD impact on rice varieties with different stomatal size and densities. (a-d) Plots relating to Fig.6 covering IR-64 and *OsEPF1oe* plants (a) Relative humidity (RH) and (b) leaf VPD and Sathi, UCP-188, Bharathy and C-7306 (c) RH and (d) leaf VPD. (e-l) Fixed RH at 55% and temperature at 39 ˚C experiment comparing Sathi, UCP-188, Bharathy and C7306 showing (e) RH, (f) chamber temperature (g), leaf VPD, (h) photosynthesis (*A*), (i) stomatal conductance ($g_{sw}$), (j) intrinsic water-use efficiency ($A/g_{sw}$; iWUE), (k) Transpiration

(*E*) and (i) Tleaf energy balance (EB). Different letters in e-l indicate a significant difference between the means (Two-way ANOVA, Tukey HSD test, $P < 0.05$). Grey diamonds represent means. a-d, $n = 7\text{-}8$ plants. e-l, $n = 5\text{-}6$ plants.
